# Recurrence-Based Information Processing in Gene Regulatory Networks

**DOI:** 10.1101/010124

**Authors:** Marçal Gabalda-Sagarra, Lucas Carey, Jordi Garcia-Ojalvo

## Abstract

Cellular information processing is generally attributed to the complex networks of genes and proteins that regulate cell behavior. It is still unclear, however, what are the main features of those networks that allow a cell to encode and interpret its ever changing environment. Here we address this question by studying the computational capabilities of the transcriptional regulatory networks of five evolutionary distant organisms. We identify in all cases a cyclic recurrent structure, formed by a small core of genes, that is essential for dynamical encoding and information integration. The recent history of the cell is encoded by the transient dynamics of this recurrent reservoir of nodes, while the rest of the network forms a readout layer devoted to decode and interpret the high-dimensional dynamical state of the recurrent core. This separation of roles allows for the integration of temporal information, while facilitating the learning of new environmental conditions and preventing catastrophic interference between those new inputs and the previously stored information. This resembles the reservoir-computing paradigm recently proposed in computational neuroscience and machine learning. Our results reveal that gene regulatory networks act as echo-state networks that perform optimally in standard memory-demanding tasks, and confirms that most of their memory resides in the recurrent reservoir. We also show that the readout layer can learn to decode the information stored in the reservoir via standard evolutionary strategies. Our work thus suggests that recurrent dynamics is a key element for the processing of complex time-dependent information by cells.

**Summary:** Cells must monitor the dynamics of their environment continuously, in order to adapt to present conditions and anticipate future changes. But anticipation requires processing temporal information, which in turn requires memory. Here we propose that cells can perform such dynamical information processing via the reservoir computing paradigm. According to this concept, a structure with recurrent (cyclic) paths, known as the reservoir, stores in its dynamics a record of the cell’s recent history. A much simpler feedforward structure then reads and decodes that information. We show that the transcriptional gene regulatory networks of five evolutionary distant organisms are organized in this manner, allowing them to store complex time-dependent signals entering the cell in a biologically realistic manner.

## Introduction

The survival of any cell, either as an individual entity or as part of a multicellular organism, depends on its capacity to respond to changes in the environment. In a wide variety of situations such as stress responses, morphogen-driven embryogenesis, immune responses, and metabolic adaptations to varying energy sources, cells need to sense multiple signals in their surroundings, integrate them, and activate an adequate response. Orchestrating the best possible response with the right intensity is crucial, but so is doing it at the right moment and promptly enough. The importance of timing and speed implies that cells able to anticipate changes in the environment have a critical advantage.

Although many changes in the environment are stochastic from the point of view of a cell, many others are predictable. In many cases, the likelihood of future events is encoded by the recent history of the cell's environment. In these cases, the ability to use the temporal character of the information is clearly beneficial. Periodic changes in the environment, for example, can be anticipated through molecular oscillators or cellular clocks, as seen in the way most organisms on earth anticipate daily light-dark cycles, including relatively simple cyanobacteria (Golden et al., 1997; Mori and Johnson, 2001). Another example is given by groups of events that tend to occur together or in a specific order. This kind of association, for instance, allows the bacterium *Escherichia coli* to prepare for oxygen depletion when it senses an increase in temperature, an indication that it has been ingested by a mammal (Tagkopoulos et al., 2008). Similarly, enterobacteria anticipate sequential changes in sugars as they pass through the intestinal tract, and yeast cells expect a specific sequence of stresses during alcoholic fermentation (Mitchell et al., 2009). Pathogenic microbes are also known to detect variations in their environment to anticipate changes in the host-pathogen interaction cycle (Rodaki et al., 2009; Schild et al., 2007). Furthermore, experimental evolution studies have shown that predictive environmental sensing can evolve in relatively short periods of time in a laboratory setting (Dhar et al., 2013).

Beyond the ability to associate concurrent events, recent studies have shown that microbes exhibit both short-and long-term memory. The stress response of *Bacillus subtilis*, for example, depends not only on the condition in which it is currently growing but also on past growth conditions (Wolf et al., 2008). However, the way in which this record of previous history— i.e. memory– is integrated and stored in cells is not yet fully understood. Knowledge of the conceptual limits of this cellular memory is also scarce. Since memory is a key limitation to recognizing temporal structures, the prediction capabilities of cells remain to be delimited as well.

While the passive prediction mechanisms of cells usually involve small circuits with only a handful of biomolecules (such as in genetic clocks), cellular adaptability relies in general on a complex network of interactions between genes and proteins, frequently at the level of transcriptional regulation (Lee et al., 2009; Martínez-Antonio and Collado-Vides, 2003). Here we hypothesize that the *global* structure of this network determines how memory is encoded. Specifically, we aim to establish how gene regulatory networks integrate complex inputs, and especially how they process time-varying information. By analyzing different organisms, we propose that gene regulatory networks encode temporal information in a state-dependent manner: the recent history of the cell is encoded in the complex transient dynamics of the network, through the interaction between its internal state (which depends on its recent past) and the external inputs that are currently being received by the system. In the field of neural networks, this strategy has come to be known as *reservoir computing* (encompassing the concepts of *echo-state network* from machine learning (Jaeger, 2001b) and *liquid-state machine* from computational neuroscience (Maass et al., 2002)).

Reservoir computing is a functional network paradigm that allows processing of temporal information while featuring a very efficient learning process. Its key characteristic is that it separates memory encoding and prediction in two different network substructures (Fig. 1). The first substructure, known as the reservoir, contains recurrent connections (i.e. cyclic paths) and encodes information by projecting the stimulus nonlinearly into a high-dimensional space (Buonomano and Maass, 2009). The recurrent network of the reservoir allows it to retain information for a certain time, providing fading memory to the system. The second substructure, the readout layer, is a feedforward structure (i.e. a directed acyclic graph) placed downstream of the reservoir. This readout layer uses the history record encoded in the state of the reservoir to make a prediction or classification. Feedforward structures, lacking cyclic paths, are much easier to train (i.e. to adapt the strength of the interactions to produce the expected dynamics). This separation of roles allows the training process to be focused solely on the readout, giving this method the computational power of a recurrent network combined with the ease of training of a feedforward architecture (Buonomano and Maass, 2009). Furthermore, by adding independent readouts the system avoids catastrophic interference, i.e. it can incorporate additional tasks without interfering with the existing ones (Lukoševičius and Jaeger, 2009).

**Figure 1:**
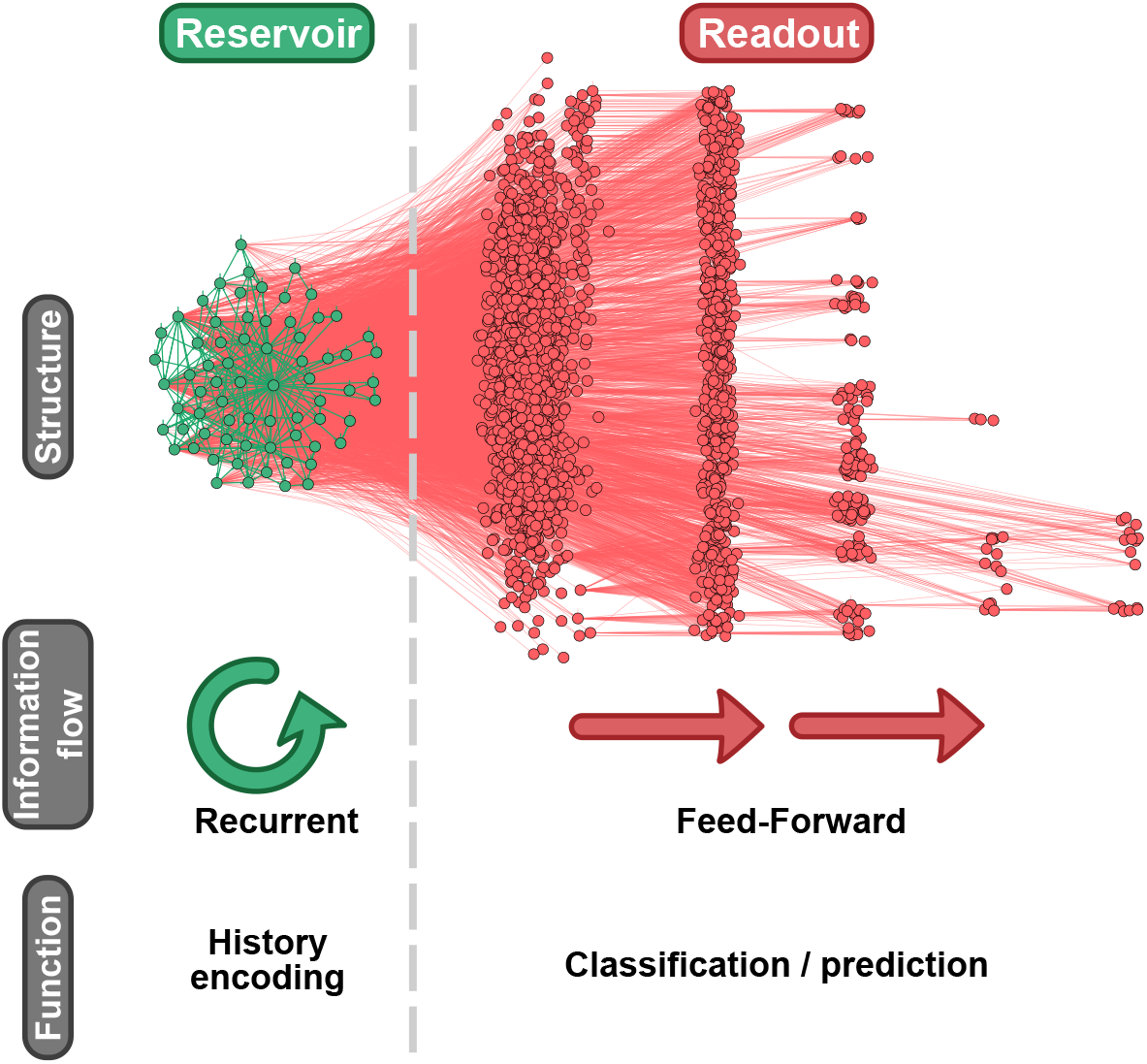
Structural and functional organization of reservoir computing. The reservoir (green) is a subgraph with cyclic paths that can maintain a record of the recent history in its dynamics. The readout (red) is a directed acyclic subgraph that reads the information encoded in the reservoir state to perform a given task. The structures shown correspond to the *Escherichia coli* network (see *Results).* The nodes in the readout are grouped by the length of the longest path reaching them from the reservoir.

In this study we propose that transcriptional networks can operate according to the reservoir computing paradigm. To do so, we first analyze the topology of the gene regulatory networks of five different, evolutionary distant organisms, and examine how efficient these networks are at encoding and processing time-varying signals. Next we show that this capability can be attributed to the reservoir-like structures found in the networks. We then consider biologically realistic inputs, in the form of different types of stress signals, and investigate how the information arriving through the corresponding pathways can be stored in the reservoirs. Finally, we show that the readout layer can be trained in a biologically realistic manner through evolutionary processes.

## Results

### Network structure

We analysed the transcriptional networks of five evolutionary distant organisms: *Bacillus subtilis, Escherichia coli, Saccharomyces cerevisiae, Drosophila melanogaster,* and *Homo sapiens.* The regulatory interaction data was obtained from different publicly available databases and publications (see *Methods* section). We limited ourselves to gene regulatory networks because in other types of cellular networks (such as protein-protein interaction networks) the directionality of the interactions, and thus of the information flow, is not so well documented. Additionally, the degree distributions of all the networks show that they have a non-trivial structure, resembling in most of the cases a scale-free architecture (Fig. S1, see also top half of Table 1 for other network descriptors).

**Table 1:** General properties of the gene regulatory networks and their recurrent cores.

|  | Nodes | Edges | Self loops | Mean degree |
| --- | --- | --- | --- | --- |
| **Whole graph** |  |  |  |  |
| *B. subtilis* | 886 | 1358 | 49 | 3.06 |
| *E. coli* | 3236 | 8366 | 126 | 5.17 |
| *S. cerevisiae* | 6725 | 201972 | 197 | 60.06 |
| *D. melanogaster* | 9432 | 231174 | 0 | 49.01 |
| *H. sapiens* | 16354 | 163271 | 28 | 19.96 |
| **Recurrent core** |  |  |  |  |
| *B. subtilis* | 13 | 30 | 7 | 4.61 |
| *E. coli* | 70 | 317 | 55 | 9.05 |
| *S. cerevisiae* | 289 | 9046 | 195 | 62.60 |
| *D. melanogaster* | 486 | 23470 | 0 | 96.58 |
| *H. sapiens* | 207 | 1434 | 26 | 13.85 |

Despite the complexity and large size of the networks, only the subgraphs containing recurrent connections are relevant for information processing according to the reservoir computing paradigm (Rodan and Tino, 2011). To identify these substructures, we pruned the networks by eliminating all the strictly feedforward nodes (see *Methods* section). The pruning process consists in iteratively removing all the nodes with either no output or no input connections until no more nodes can be removed. This procedure leads to a single main recurrent structure in each network, which will be referred to in what follows as the *core* or *reservoir* of the network (bottom half of Table 1). As can be seen in Fig. 2, the recurrent cores in all five cases are much smaller, in terms of number of genes involved, than the corresponding whole network (see also Table 2).

**Figure 2:**
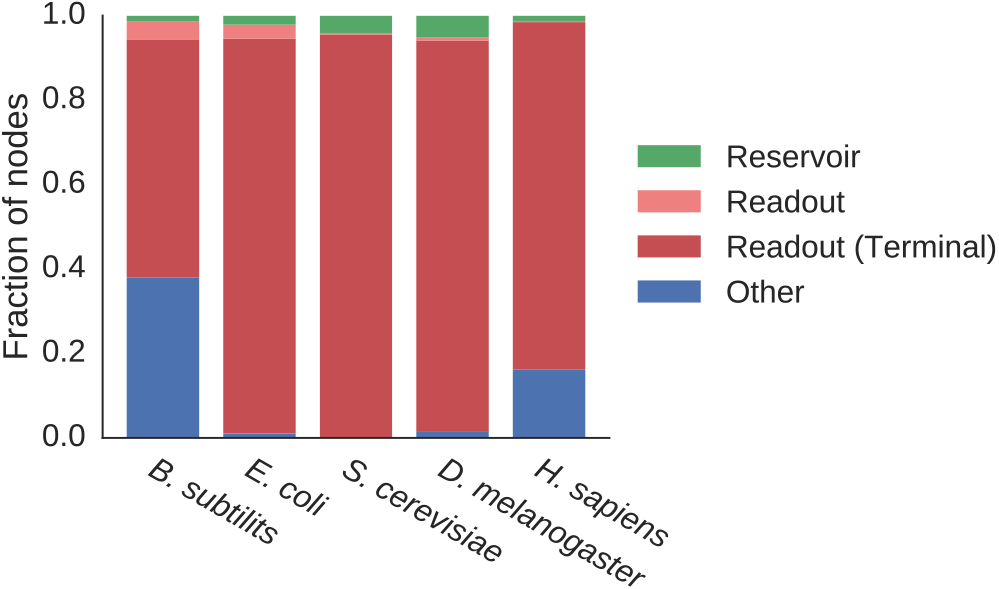
Relative sizes of functional groups for each network. The fraction of the total number of nodes that belong to each network substructure are shown. Reservoir nodes are the ones left over after network pruning. The nodes placed downstream of the reservoir are assigned to the readout structure, distinguishing between terminal nodes, which have zero out-degree, and the rest. Finally, all nodes that do not fall in any of the previous groups are counted as ‘others’.

In order to establish where each small subgraph of nodes is located within its network, we also show in Fig. 2 the fraction of genes located downstream of the reservoir (forming what we call the *readout,* following the organization depicted in Fig. 1). It can be observed that the vast majority of nodes are placed downstream of the recurrent core (the nodes labeled *other* are either upstream of the reservoir or fully isolated)^1^. The obvious implication of the location of the core is that most of the network is affected by its dynamics. It is worth noting again that by definition there are no recurrences outside the reservoir, and thus none of the readout nodes can affect back the reservoir. Furthermore, Fig. 2 also shows that a very large proportion of the readout nodes are terminal (i.e. nodes with no output connections). That limits the potential complexity of the readout topology, and thus its ability to process information, giving an even more central role to the recurrent core or reservoir, as we show below.

**Table 2:** Number of nodes in the different network substructures.

|  | Total | Reservoir | Readout (terminal) | Other |
| --- | --- | --- | --- | --- |
| *B. subtilis* | 886 | 13 | 537 (500) | 336 |
| *E. coli* | 3236 | 70 | 3133 (3025) | 33 |
| *S. cerevisiae* | 6725 | 289 | 6436 (6419) | 0 |
| *D. melanogaster* | 9432 | 486 | 8795 (8721) | 151 |
| *H. sapiens* | 16354 | 207 | 13497 (13449) | 2650 |

### Encoding ability

Next, we inquired if these recurrent cores are able to encode temporal information in their dynamics. To do so we confronted them to the 10th order Nonlinear Auto-Regressive Moving Average (NARMA) task, a memory-demanding benchmark commonly used in the reservoir computing context (Appeltant et al., 2011) (see *Methods* section). To test if the dynamics of the network cores can represent the recent history, the network was simulated with simplified dynamics and a time-varying random input (*zt* in Fig. 3) was applied to it. Then, an *ad hoc* readout node was trained (i.e. the readout weights W^out^ are adjusted) to reproduce the output *yt* of the 10th order NARMA system using only the instantaneous state of the network (Fig. 3). The challenge is that the output of the 10th order NARMA task depends on the input and output values of the last 10 time steps. This information about the past must be encoded in the reservoir state for the output *ỹ*_*t*_ of the readout node to be able to accurately model the NARMA system. Fig. 4 shows a representative time trace of the input signal *zt,* the actual NARMA system output *ỹ*_*t*_, and the reconstruction *yt* obtained with each of the biological networks. The figure shows that the reconstructed output mimics closely the expected output for large enough reservoir sizes (bottom four rows, see Table 2 for core sizes).

**Figure 3:**
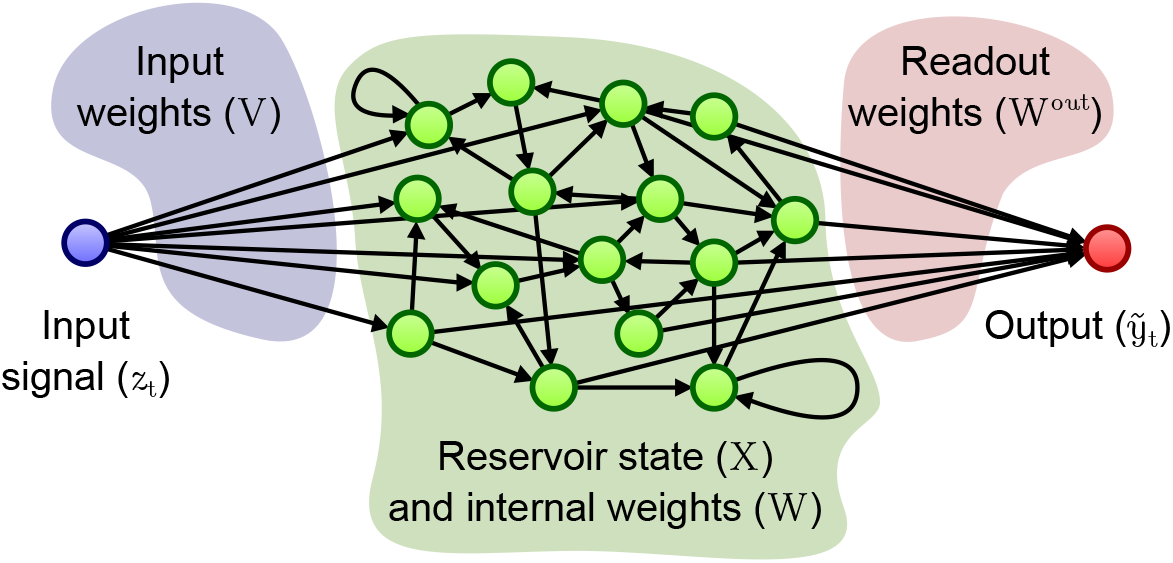
Setup to test the memory of a network. A reservoir is built with a connectivity matrix *W* extracted from the topology of the biological network. An input signal *z*_*t*_ is applied to the nodes of the reservoir with different strengths, defined by the input weight vector *V*. Then, one or more readout nodes compute a weighted sum of the state of the reservoir X. The weight vector *W*^out^ is tuned so that the output *ỹ*_*t*_ of the readout approximates a target output signal *y*_*t*_.

**Figure 4:**
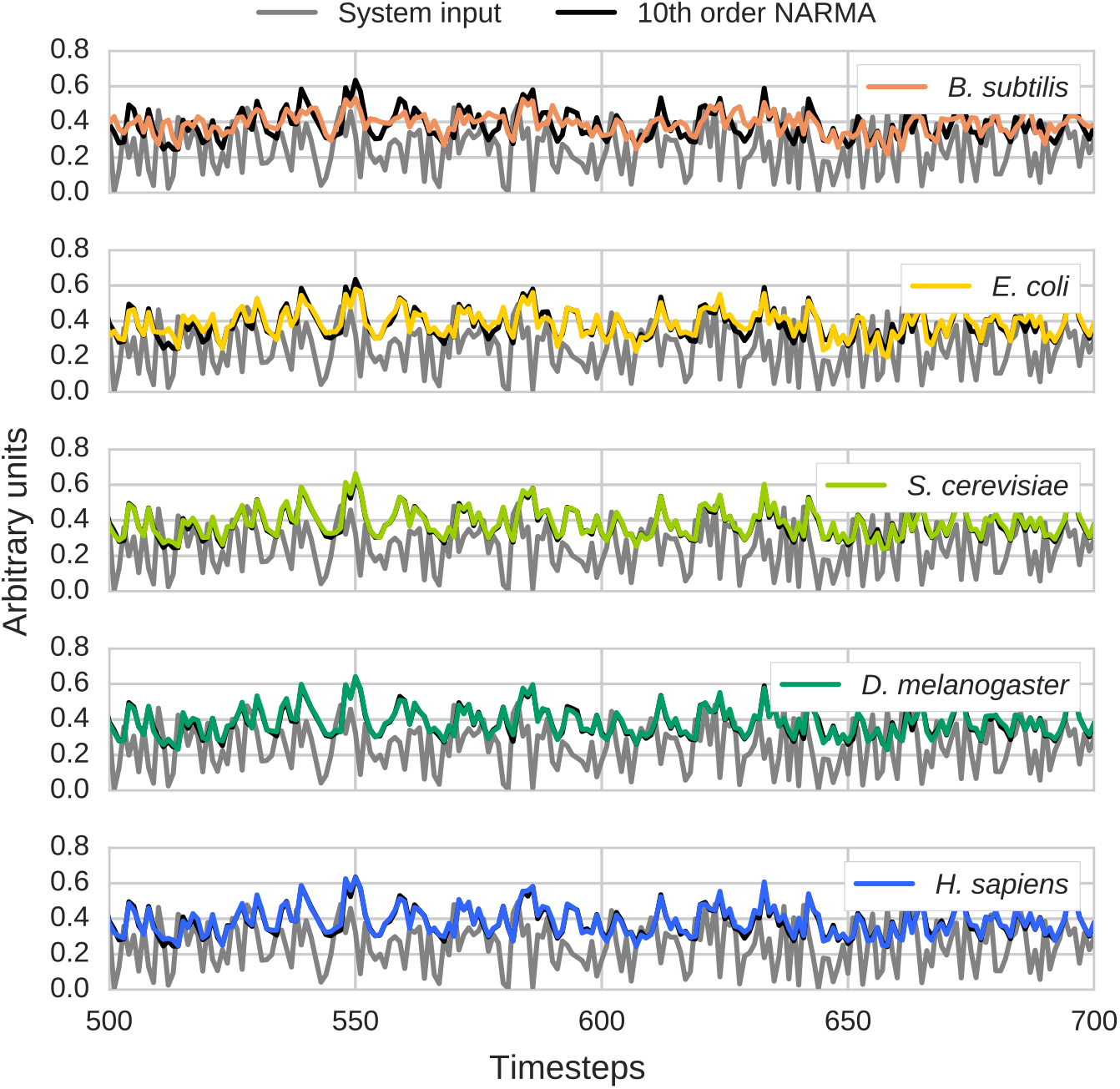
Representative time series of the test phase of a 10th order NARMA task. For each biological network studied, pruning was used to build a reservoir core, and a readout node was trained to reconstruct the output of the 10th order NARMA system, using the state of this reservoir. Gray lines represent the random input of the system, black lines the actual output of the NARMA function, and colored lines the output reconstructed by the readout node in each case.

We next compared the performance of our biological networks with the *de facto* standard topologies in the reservoir computing literature. To quantify this performance we used the Normalized Root Mean Squared Error (NRMSE) between the reconstructed and the expected output signals (see *Methods* section). Fig. 5 shows the median NRMSEs achieved by our reservoirs, and compares them with control topologies of diverse core sizes. The control networks take the form of random (Erdös-Renyi) echo-state networks for which the network density (dESN) or mean degree (kESN) are kept constant (and equal to their biological counterparts) as the network size varies. We also include results for simple cycle reservoirs (SCRs), linear cyclic reservoirs with the minimum recurrence that allows them to operate as echo-state networks (Rodan and Tino, 2011). As the figure shows, all the biological cores perform as well as the random dESN and kESN control networks of the same size, and for large enough core sizes (*S. cerevisiae, D. melanogaster, H. sapiens*), the performance is much better than the corresponding SCR. In fact, differences between SCR and both the biological networks and ESN variants increase with size within the interval analysed. Results also suggest that the different performance of each GRN is related with their size. In this regard it is worth noting that despite the fact that the number of edges in the control networks scales linearly with the size for kESNs and quadratically for dESNs, they show similar performance to each other for all the range of sizes. That discards any major effect of the number of edges in the performance of the reservoir in these conditions.

**Figure 5:**
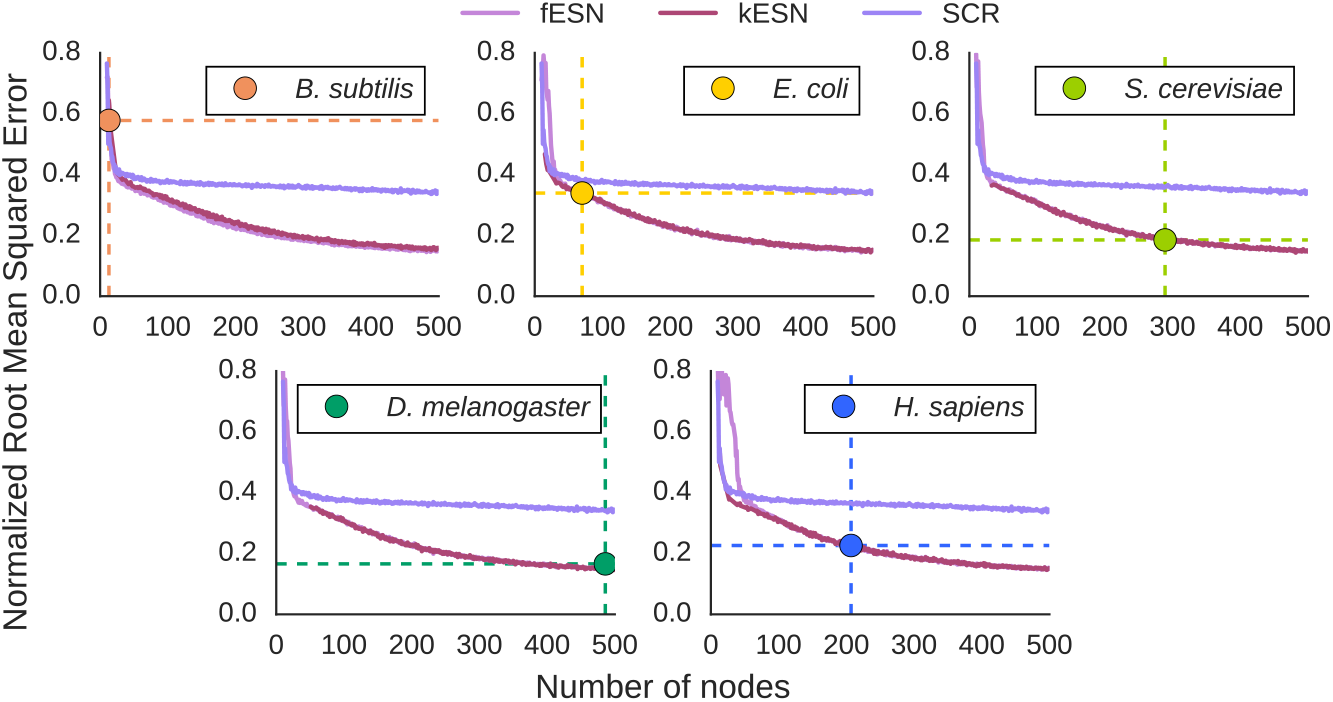
Performance of the biological reservoirs compared with control topologies. Performance is evaluated with the Normalized Root Mean Squared Error (NRSME) between expected and reconstructed outputs. The NRMSE value shown for each biological network topology corresponds to the median of 10000 trials (with edge weights and data series randomization). The values plotted for each control network (dESN, kESN and SCR) correspond to the median value of 100 trials for each network size from 10 to 500 nodes. In each case, dESN and kESN are produced keeping the network density and mean degree, respectively, of the biological core with which they are compared (see *Methods* section).

To quantify the amount of temporal information that our networks can store, we computed their critical memory capacity (maximum number of past time steps that can be recovered with a given accuracy). For each gene regulatory network, we applied this test to three subnetworks: the recurrent reservoir, the readout, and the largest connected component (which contains the first two). The results shown in Fig. 6 confirm that most of the ability of our networks to encode history is provided by their reservoirs (the green and blue bars in the figure have very similar heights in all cases). In contrast, the readouts have much lower memory capacities, in spite of being much larger in size than the reservoirs, see Fig. 2. The critical memory capacities of the readouts are mainly determined by length of the longest path (Fig S2), as expected from their feedforward structure. These results confirm that the recurrent cores are responsible for most of the capacity to dynamically store temporal information of the gene regulatory networks.

**Figure 6:**
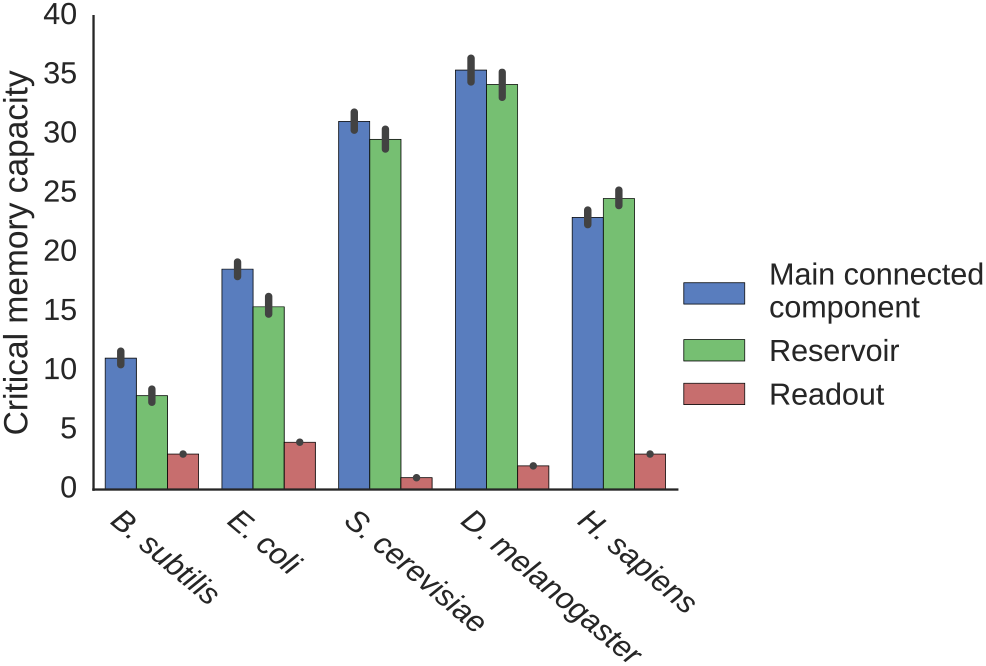
Critical memory capacity of the different network substructures. Results are shown for the reservoir (green) and readout (red) substructures, and for the largest connected component—which comprises the other two— (blue), for each of the gene regulatory networks analyzed. Median values are shown (*n* = 30 trials). The error bars indicate the 98% confidence interval (CI) computed by bootstrapping.

We have considered so far that information is introduced in the reservoirs via randomly selected nodes. However, realistic biological inputs act upon specific sets of nodes. To determine whether the reservoir computing paradigm holds in the presence of such realistic inputs, we identified nodes from the reservoir that belonged to particular stress-response pathways, and studied the effect of the corresponding stresses. We worked specifically with the E. coli reservoir in order to keep a balance between methodological tractability and network performance. For each stress type, each reservoir node was scored depending on the level of evidence (according to the literature cited in the EcoCyc database) supporting that the stress signal acts upon it. Table 3 lists the number of nodes that are considered to receive information from every stress with a confidence score equal or larger than a given threshold (see Dataset 2 for a detailed list).

We next subjected the *E. coli* reservoir to a variation of the NARMA test, in which the stress signals act solely upon the input genes selected above. Figure 7 shows the resulting NRMSE as a function of the number of input nodes for the different stresses (encoded with distinct colors), and for all confidence thresholds (which correspond to different sizes, according to Table 3). As a control, the figure also shows the NRSME obtained applying the input signal to random sets of nodes (gray line). The most obvious conclusion from these results is that biologically realistic inputs are as efficient as randomly selected nodes at encoding information in the reservoir. Also, the precision of the system increases monotonically with the size of the input set, and eventually saturates. Besides, it is noteworthy that although the different stress signaling pathways affect different sets of nodes, their ability to introduce information in the system is comparable. This highlights the fact that memory is encoded in the network in a delocalized manner, without depending on specialized circuits or structures.

**Table 3:**
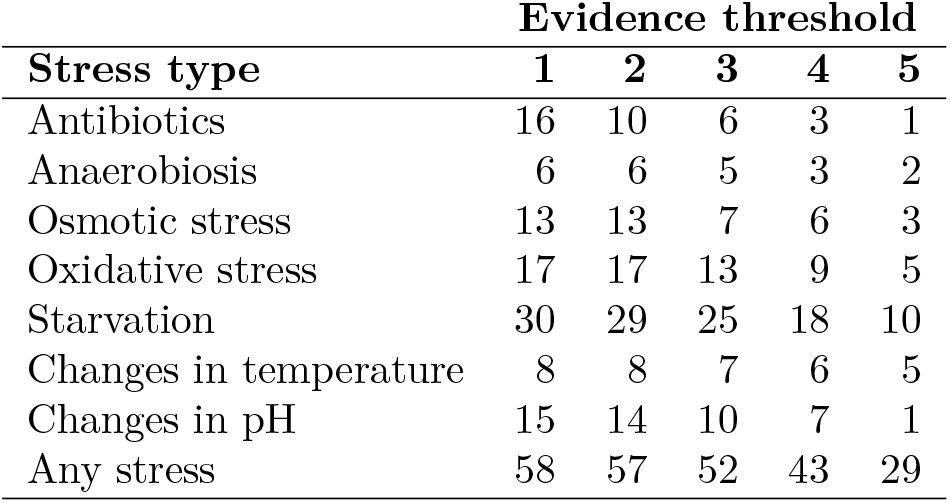
Number of nodes affected by stress type for each evidence threshold.

| Stress type | Evidence threshold |  |  |  |  |
| --- | --- | --- | --- | --- | --- |
|  | 1 | 2 | 3 | 4 | 5 |
| Antibiotics | 16 | 10 | 6 | 3 | 1 |
| Anaerobiosis | 6 | 6 | 5 | 3 | 2 |
| Osmotic stress | 13 | 13 | 7 | 6 | 3 |
| Oxidative stress | 17 | 17 | 13 | 9 | 5 |
| Starvation | 30 | 29 | 25 | 18 | 10 |
| Changes in temperature | 8 | 8 | 7 | 6 | 5 |
| Changes in pH | 15 | 14 | 10 | 7 | 1 |
| Any stress | 58 | 57 | 52 | 43 | 29 |

**Figure 7:**
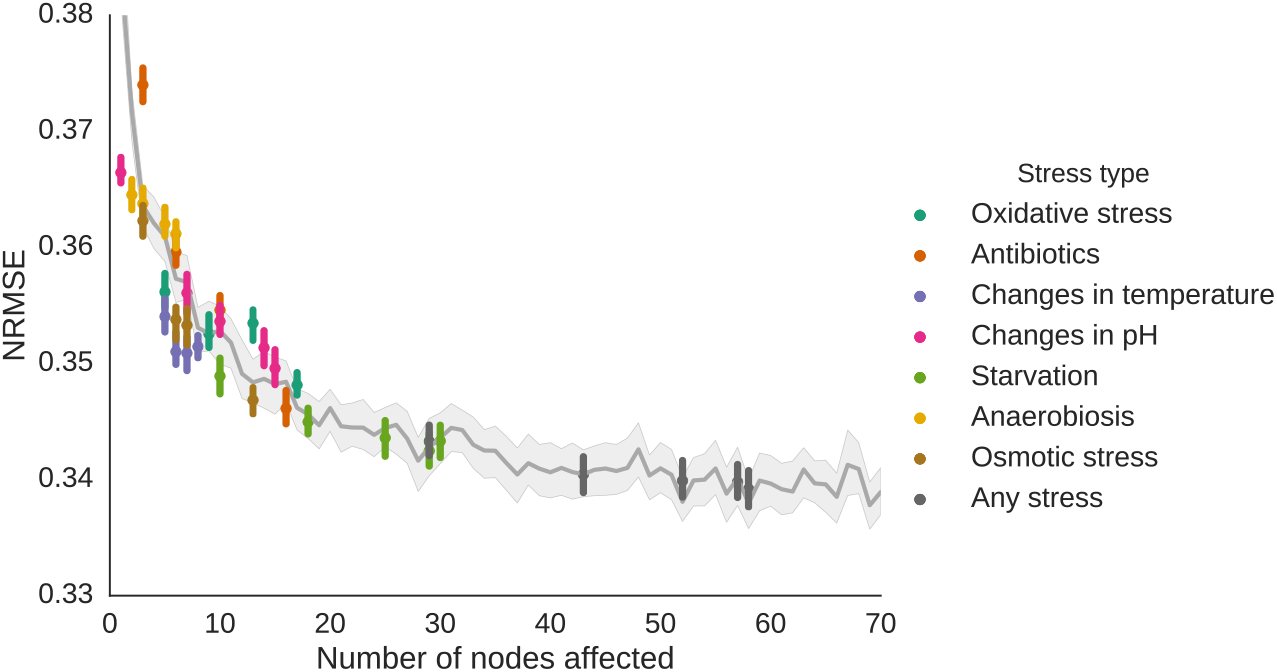
Efficiency of signaling pathways feeding an input signal to the reservoir. Median NRMSE obtained in the NARMA test using as input nodes those genes affected by signaling pathways that react to different stress types. The gray line marks the NRMSE obtained when applying the input to a random set of nodes of a given size. The median values are shown of 1000 replicates for each random input size and 2000 for each biological input set. Error bars and shaded area indicate the 98% CI computed by bootstrapping.

Finally, we tested whether biological processes can shape a readout structure that can use the temporal information encoded in the dynamics of a reservoir. Specifically, we examined if a readout can be evolved in a situation where the information about recent events gives a evolutionary advantage. For that purpose, we simulated an evolutionary process using the covariance matrix adaptation evolutionary strategy (CMA-ES) (Jiang et al., 2008). Using the *E. coli* reservoir, a population of single-node readouts was let to evolve with a selective pressure to predict the 10th order NARMA system. Figure 8 features a representative instance of such evolutionary processes (green), compared with the behavior of a standard ridge regression training method (purple). The performance of both types of readouts were practically indistinguishable after 2000 generations, indicating that evolution can, indeed, tune a readout structure to read the temporal information stored in a reservoir.

## Discussion

In the present study we propose a new paradigm to understand how cellular regulatory networks can store and process temporal information. Specifically, we suggest that these networks can function as reservoir computing systems. A division of labor allows to separate the processes of memory encoding and decision making in two distinct regions of the network. The first region, the reservoir, has recurrences—i. e. *cyclic* paths— that give it a *fading memory* property so that it can efficiently encode recent history. The latter region, the readout, has a feedforward or acyclic structure and uses the information it receives from the reservoir to make a classification or prediction. This separation of roles allows the system to process temporal information while still being very efficient when learning new tasks (Buonomano and Maass, 2009).

The results of analyzing gene regulatory networks of five evolutionary distant organisms support this hypothesis. First, the topology of all five networks matches the structural characteristics of a reservoir computing system. Second, we show that these loosely defined reservoir structures are able to encode in their dynamics an amount of temporal information that is non-trivial given their sizes. As a matter of fact, for all networks the reservoir is one to two orders of magnitude smaller than the readout, and yet its critical memory capacity is up to around 30 times higher. Moreover, in the case of *E. coli*, biological signals relevant for the cell (specifically, physiological stresses) arriving at the reservoirs at different locations are encoded with similar performance, indicating that the ability to store temporal information is distributed in the reservoir. This independence of the specific entry point of the information in the system confers robustness to failure: while some of the input streams may get compromised, it is unlikely that a large group of them would fail simultaneously. Finally, we show that evolution can produce readout structures that are able to decode the state of a reservoir, as long as temporal information provides a selective advantage.

Non-recurrent gene network architectures have been proposed in the past as mechanisms of information integration and storage (Bray, 1995; Scheres and Van Der Putten, 2017), associative learning (McGregor et al., 2012; Sorek et al., 2013), and cellular decision making (Bates et al., 2015; Filicheva et al., 2016). However, processing of time-dependent information requires recurrent topologies such as the ones investigated in this paper. The nonconventional computation framework proposed here also implies that the integration of information is distributed across the network in large and diffuse structures with well-defined functional roles. A similar connection between gene regulatory networks and reservoir computing systems was hinted by Jones *et al* (Jones et al., 2007). However, that study uses as putative reservoir a gene regulatory network that does not include any recurrences other than self-regulations of some scarcely interconnected nodes, and the system is tested with a task that does not require memory.

Even though it is clear that cells benefit from anticipating the environment, we know no example yet of a single cellular system that processes complex temporal information in nature. The most studied types of anticipation, involving periodic (Golden et al., 1997; Mori and Johnson, 2001) and sequential (Mitchell et al., 2009; Tagkopoulos et al., 2008) events are mechanistically fairly simple. However, *ad hoc* experiments have shown that *Physarum polycephalum*, also known as slime mold, and *Plasmodium cudatum* can learn very efficiently new temporal structures. Slime mold, in particular, can anticipate a shock after experiencing a single series of three ten-minute low-temperature shocks at one-hour intervals. Moreover, if the organism experiences a new shock several hours later, it pre-emptively reacts to the two missing following shocks (Saigusa et al., 2008). Similarly, *P. cudatum* can learn that an electric shock follows an innocuous vibratory or luminous stimulus (Armus et al., 2006; Hennessey et al., 1979). All these results hint at capabilities to learn temporal structures larger than what can be easily explained with current models.

We propose that cells can process temporal information and anticipate their environment by using their regulatory networks as computational reservoirs. To that end, here we explored the potential of transcriptional networks to encode the recent history of cells, but other regulatory networks such as protein-protein interaction or metabolic networks may play a similar role. The combination of the different timescales (minutes or seconds) and learning mechanisms (evolution, chromatin regulation for transcriptional reservoirs, or expression regulation for post-transcriptional reservoirs) could give rise to much richer behaviours.

In our study, the dynamics of the networks have been largely simplified with a formalism used for neural networks. The real dynamics, with nonlinear interactions and different time scales for each gene, would add more complexity to the network behaviour and increase the memory of the system (Büsing et al., 2010; Dambre et al., 2012; Tanaka et al., 2016). Furthermore, the interaction of layers of regulatory networks with different time scales—e.g. transcriptional, protein protein interaction or metabolic networks— also could increase the memory capacity of the system (Dambre et al., 2012; Gallicchio and Micheli, 2016). Far from invalidating our results, our simplification of the dynamics makes our tests more stringent. Additionally, the NARMA task is known to be highly demanding, as it requires a significant level of precision in the results. Probably life does not need to be as precise.

## Methods

### Cellular regulatory networks

We used published transcriptional regulatory interactions to build our gene regulatory networks. Data for *Bacillus subtilis* was obtained from DBTBS (Sierro et al., 2008). Data for Escherichia coli was extracted from EcoCyc (Keseler et al., 2011), including the sigma factors as transcription factors. Data for Saccharomyces cerevisiae was obtained from YEASTRACT (Teixeira et al., 2014). The gene regulatory network for Drosophila melanogaster was obtained from the modENCODE initiative (Roy et al., 2010). Finally, data for Homo sapiens was extracted from the ENCODE project (Gerstein et al., 2012).

### Network pruning

The networks were simplified to a minimal recursive subgraph, i.e. a subgraph containing only the nodes and edges that form cyclic paths and the nodes that interconnect them. To do so we *pruned* the networks by iteratively removing any node that had either in-degree or out-degree equal to zero, until no more nodes could be removed.

### Simulation of network dynamics

The dynamics of the gene regulatory networks were simulated using a discrete time updating rule defined as

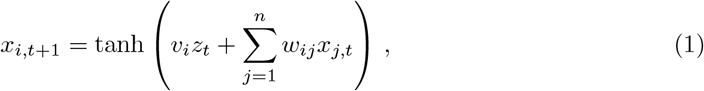

where we follow the notation shown in Fig. 3: *z_t_* is the system input at time *t, xi, t* is the state of the *i* node of the reservoir at time t (corresponding to the elements of the reservoir state vector *X*), *n* is the number of nodes in the reservoir, *w_ij_* is the weight of the link from the *j*th to the *i*th node (representing the elements of the weighted adjacency matrix *W*), and vi is the weight of the link from the input to the ith node (corresponding to the elements of the input weight vector *V*). The values of vi are randomly chosen to be either -0.05 or 0.05. At the same time, the values of wij are real numbers drawn from a uniform distribution between -1 and 1 if the link exists, and 0 otherwise (if the sign of the interaction, i.e. activation vs repression, is known, the sign of wij is set accordingly). Additionally, the W matrix was normalized to have a spectral radius of 0.9 to assure the echo-state property (Jaeger, 2001b; Lukoševičius and Jaeger, 2009).

### NARMA task

The Nonlinear Auto-Regressive Moving Average (NARMA) task has been widely used as a memory test in the context of reservoir computing (Jaeger, 2002). It consists in training a network to model the output of the 10th order NARMA system (Atiya and Parlos, 2000), a discrete time system where the input values s(t) are drawn from the uniform distribution U(0, 0.5) and the output y(t) is defined by

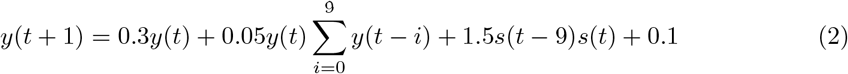

For the task, we simulated a network with the studied topology with a single input node feeding the *s*(*t*) series in the system. A readout node was then trained via ridge regression (see below) to model the *y*(*t*) series. For each realization a NARMA series of 10000 steps was generated, using 9000 of them for the training phase and 1000 to test the performance. The evaluation of the NARMA modeling was done using the normalized root mean squared error measure, defined as

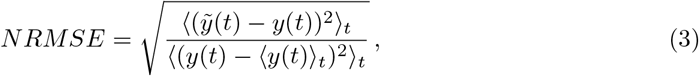

where 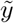(*t*) is the output predicted by readout, *y*(*t*) is the output of the actual NARMA system, and <·>_*t*_ indicates the mean over time.

The following topologies were used as controls:

- Echo State Network – fixed mean degree (kESN): Erdös-Rényi random network with the same mean degree (2 × *n*_edges_/*n*_nodes_) as the topology of the corresponding biological network.
- Echo State Network – fixed network density (dESN): Erdös-Rényi random network with the same network density—i.e. fraction of existing links over all possible ones—(*n*_edges_/*n*^2^_nodes_) as the topology of the corresponding biological network.
- Simple Cycle Reservoir (SCR): a directed circular graph, which is the simplest topology that can work as a computational reservoir (Rodan and Tino, 2011).

Note that for control networks with the same number of nodes as the problem topology, kESN is equal to dESN. This is not the case, however, when the number of nodes changes.

### Ridge regression nodes

A ridge-regression readout computes a weighted sum of the state of the nodes it receives information from (Fig. 3):

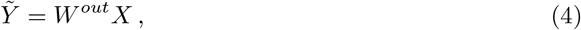

where *W^out^* is a vector of the *w_i_^out^* i weights given to the ith node by the readout, and *Ỹ* is a vector with all predicted outputs over time. A ridge regression is a type of linear regression in which the regression coefficients are obtained from

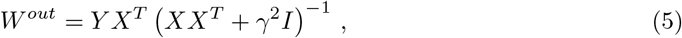

where *X^T^* is the transpose of *X*, *Y* is a matrix with all expected outputs over time, *I* is the identity matrix and γ is a regularization parameter.

Ridge regression favors regression coefficients with smaller absolute values. In doing so it introduces a certain bias, but on the other hand it also reduces the variance of the estimate. This allows estimating the parameters of a linear regression when the predictor variables are strongly correlated, making it a common readout choice in the context of reservoir computing (Wyffels et al., 2008).

### Critical memory capacity

To quantify the memory of our networks, we applied a variation of the short-term memory capacity (Boedecker et al., 2012; Jaeger, 2001a). Specifically, we simulated the network with a single input node feeding a signal *u*(*t*) drawn from a random uniform distribution between -1 and 1. Then, a ridge-regression node was trained to obtain an output 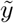(*t*) that aims to reconstruct a delayed version of the input signal *u*(*t*-*k*). The *k*-delay memory capacity (*MC_k_*) is then defined as

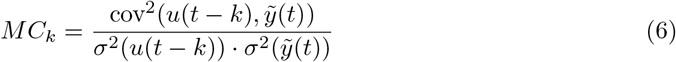

The short-term memory capacity is typically defined as 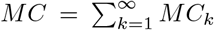, where the infinite summation is approximated by a long enough finite one (Boedecker et al., 2012; Jaeger, 2001a). The limitation, though, is that the time series needs to be orders of magnitude longer than the size of the network to ensure that lim*_k→∞_MC_k_* = 0. Otherwise, *MC_k_* will never reach 0 and *MC* will never converge. Since we dealt with fairly large networks, computing the short-term memory capacity with a reasonable precision was not feasible. As an alternative measure we defined the critical memory capacity *k\** as the maximum delay *k* that fulfills *MC_k_* > 0.5.

### Stress signal inputs

We analyzed which of the 70 genes present in the *E. coli* reservoir were known to be affected by signaling pathways that react to different stress classes. Biological stresses of seven different classes were considered, namely: presence of antibiotics, anaerobiosis, osmotic stress, oxidative stress, starvation, changes in temperature, and changes in pH. Using the annotations of the Ecocyc database (Keseler et al., 2011), we manually set a confidence score each possible stress-gene interaction. This score indicates the level of evidence supporting that the product of a given gene is affected by a signaling pathway in response to a given stress. Both post-transcriptional and transcriptional regulations were considered to determine if signaling pathway could reach a given gene, as long as they were not already included in the network structure (i.e. only transcriptional interactions coming from outside the reservoir were considered).

Using this information, an input weight vector *V* was constructed for each stress class and confidence threshold. Given a threshold, all interactions with lower score were set to zero while the others were initiated normally. Additionally, the sign of each entry was set to be positive (negative) if the interaction was known to produce an activation (repression) of the gene, and randomly set otherwise.

### Evolutionary training

Reservoir computing usually relies on ridge regression, described above, to train the readout weights. Cellular networks, however, should use biologically realistic means to perform this training. A reasonable possibility is that the readout weights are tuned through evolutionary processes. To assess this possibility, we used here an evolutionary algorithm. Starting from a first generation of random candidate weight vectors, we iteratively generate new generations by duplicating and introducing variations to the best performing solutions in the current generation.

Specifically we used the implementation of the Covariance Matrix Adaptation Evolutionary Strategy (CMA-ES) algorithm (Jiang et al., 2008) provided in the package ‘Distributed Evolutionary Algorithms in Python’ (DEAP) (Gagn, 2012). The CMA-ES algorithm learns the covariance matrix of mutations in successful individuals, so that beneficial mutations are sampled more often. This approach reduces the computational cost while preserving the biological relevance.

In Fig. 8, a single input node feeds into the *E. coli* reservoir a *s*(*t*) signal drawn from a random uniform distribution *U*(0, 0.5). Besides, the input vector *V* was defined so that the signal would only reach genes known to be affected by at least one stress type with a confidence score of 3 or more. Then, a population of readout nodes represented by their weight vectors *W^out^* was let to compete and evolve, giving a selective advantage to the ones that reproduced better the output of the NARMA system. Furthermore, for each generation in the evolutionary process a new realization of the *s*(*t*) input signal was used, recomputing the reservoir dynamics and the expected output. On the other hand, the internal weights of the reservoir *W* and the input weights *V* were kept constant for the whole simulation.

**Figure 8:**
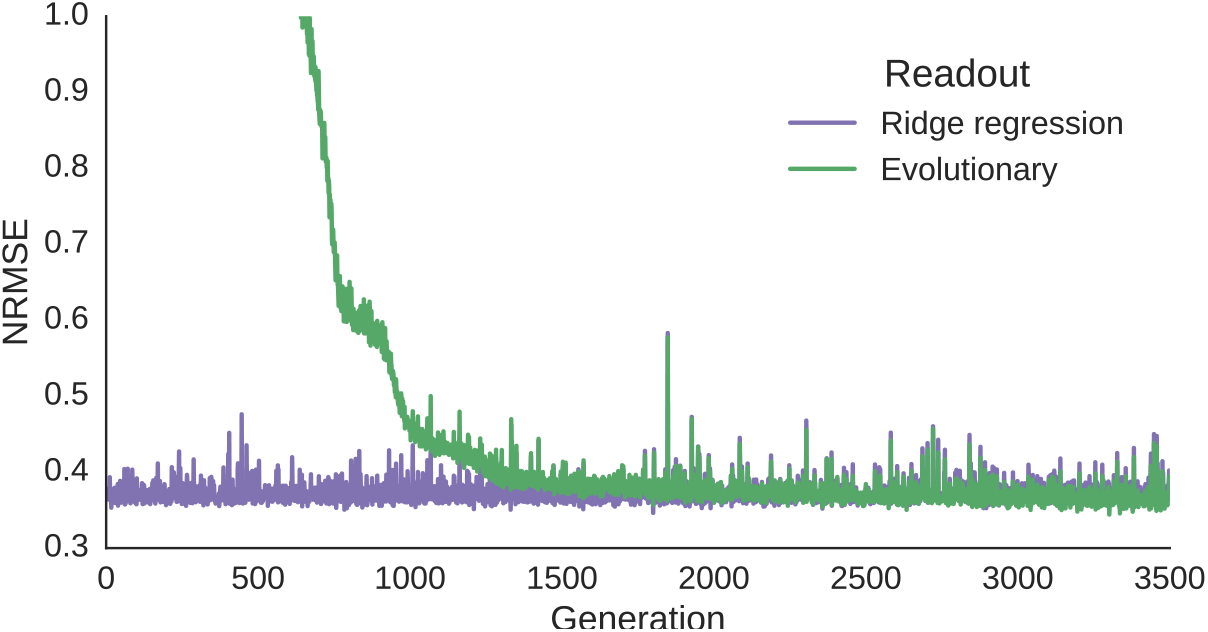
Training of a readout through an evolutionary process. NRMSE during the evolutionary training process of a readout for the *E. coli* reservoir. The weights of the reservoir node were trained using the CMA-ES (Jiang et al., 2008) algorithm to model the 10th order NARMA system Conceptually, a population of 200 candidate solutions is let to evolve while giving a selective advantage to those obtaining a lower NRMSE. The green line shows the NRMSE of the centroid of the best solutions in each generation. The purple line shows the NRMSE obtained in the same situation by a readout trained using standard ridge regression.

## Acknowledgements

We thank Rosa Martinez-Corral for useful comments. This work was supported by the Spanish Ministry of Economy and Competitiveness and FEDER (project FIS2015-66503-C3-1-P), and by the Generalitat de Catalunya (project 2014SGR0947). J.G.O. also acknowledges support from the ICREA Academia programme and from the “María de Maeztu” Programme for Units of Excellence in R&D (Spanish Ministry of Economy and Competitiveness, MDM-2014-0370).

## Supporting information

**Figure S1:**
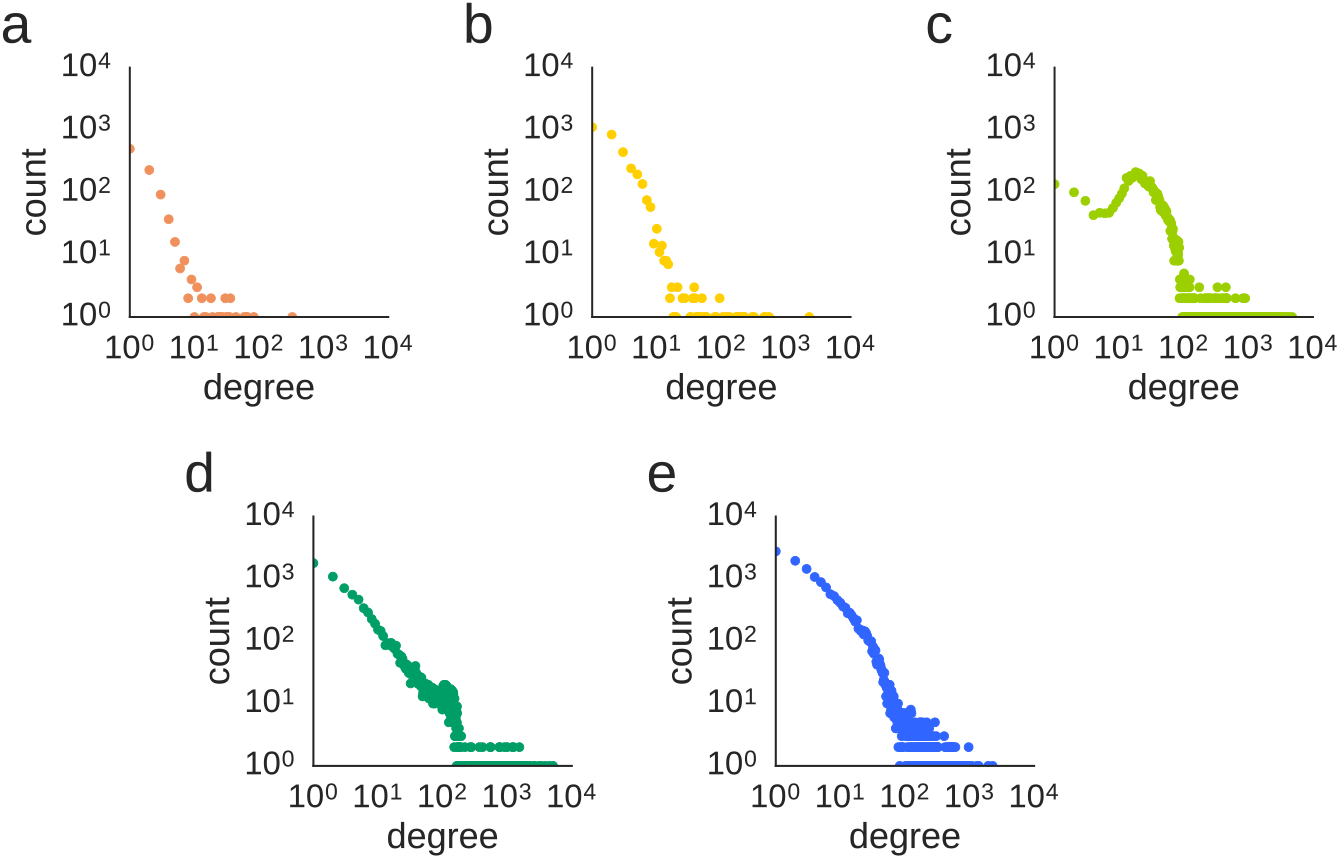
Degree distribution of the gene regulatory networks. Each plot corresponds to one of the networks: *Bacillus subtilis* (A), *Escherichia coli* (B), *Saccharomyces cerevisiae* (C), *Drosophila melanogaster* (D), and *Homo sapiens* (E). In the case of *Saccharomyces cerevisiae*, the deviation observed in the degree distribution plot (panel C) is thought to be an artefact: since most of the data in this database comes from compiling a large number of low throughput studies, nodes with lower degree can be expected to be under-represented, as studies tend to focus on genes involved in more regulatory interactions.

**Figure S2:**
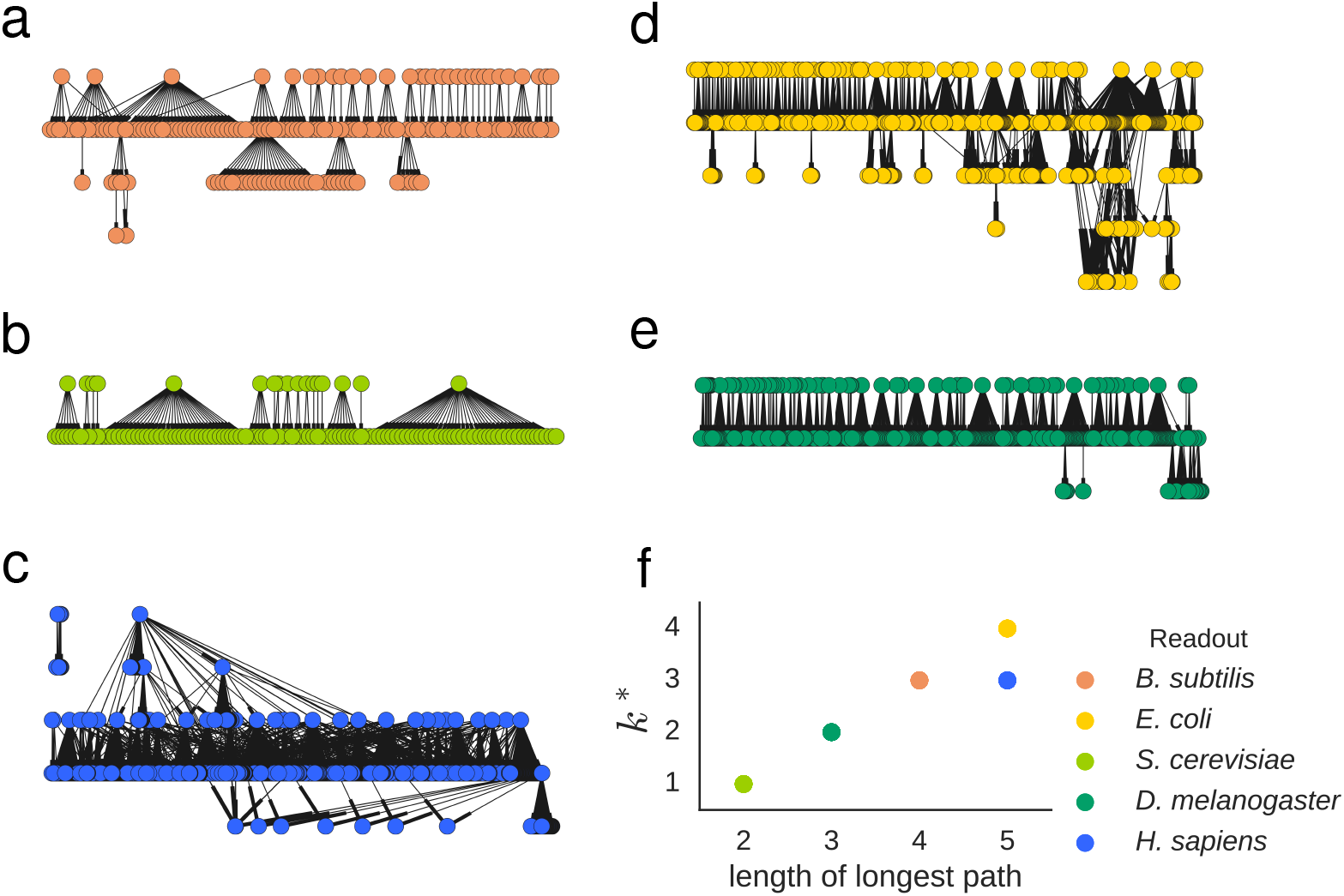
Topology and critical memory capacity of the readout structures of the gene regulatory networks. Hierarchical representation of the readout of the five gene regulatory networks: *B. subtilis* (A), *E. coli* (B), *S. cerevisiae* (C), *D. melanogaster* (D), and *H. sapiens* (E). Nodes are ordered in layers from top to bottom according to the length of the longest path reaching them from the reservoir. Panel (F) shows the relation between the length of the longest path in each of the readouts and their critical memory capacity *k\**, which measures the number of time steps in the past that can be remembered with a certain precision in the system dynamics.

## Footnotes

1 The list of genes belonging to each of the substructures for the five networks, and the connections between them, is given in Dataset 1.

